# Sex-specific disruptions in the developmental trajectory of anxiety due to prenatal cannabidiol exposure

**DOI:** 10.1101/2024.12.02.626301

**Authors:** Daniela Iezzi, Alba Cáceres-Rodríguez, Pascale Chavis, Olivier J. Manzoni

## Abstract

Many pregnant women use cannabidiol (CBD) as a natural remedy to alleviate symptoms such as nausea, insomnia, anxiety, and chronic pain. As much as 20% of pregnancies in the USA and Canada may involve the use of CBD-only products. CBD crosses the placenta and may affect fetal development, potentially leading to neuropsychiatric conditions later in life. Given the limited understanding of the effects of CBD during pregnancy, we adopted a longitudinal approach to investigate the neurodevelopmental trajectory associated with prenatal CBD exposure.

Pregnant mice were administered 3 mg/kg CBD from gestational days 5 to 18. At early adolescence, offspring displayed sex-specific behavioral changes. Females, but not males, exhibited a complex anxiety-like phenotype during the elevated plus maze task. This phenotype persisted into adulthood in the open field test and was accompanied by altered reward responsiveness. Throughout post-natal life, female offspring demonstrated heightened stretch-attend postures, a risk-assessment behavior reflecting approach-avoidance tendencies and anxiety. Finally, prenatal CBD exposure increased repetitive behaviors in adult animals of both sexes, as evidenced by the marble burying task. These results provide strong evidence of sex-specific disruptions in the developmental trajectories of anxiety associated with prenatal CBD exposure. They challenge the perception that CBD is universally safe and highlight vulnerabilities linked to gestational CBD exposure.

## Introduction

During in utero development, the brain is exquisitely sensitive to its environment. Disruptions to this development can lead to significant changes in brain structure, connectivity, and behavior. Exposure to environmental factors, such as drugs, during critical periods of development can have lasting effects on an offspring’s health, potentially resulting in neurodevelopmental and neuropsychiatric disorders. [1], [2]. These disorders, including anxiety, major depression, bipolar disorder, schizophrenia, and obsessive-compulsive disorder, affect millions of people worldwide. Increasingly, research suggests that many neuropsychiatric conditions, which may not manifest until childhood or early adulthood, originate in the fetal period. [3].

Drug use during pregnancy poses significant concerns for both the mother and the developing baby. Cannabis is the most commonly used psychoactive substance among pregnant women. [4]–[8]. Many turn to cannabis to alleviate symptoms like nausea, weight gain, and sleep difficulties [9]. The primary active compounds, Δ9-tetrahydrocannabinol (Δ9-THC) and cannabidiol (CBD), can cross the placenta [10], [11], entering the baby’s bloodstream, and can also accumulate in breast milk [12], exposing the baby after birth. This raises concerns about potential risks to the baby’s health and development [13]. Studies in humans have shown that maternal cannabis use may have long-term effects on a child’s development [for extensive review see Natasha et al.[14]]. Additionally, animal studies with synthetic cannabinoids or isolated THC indicate that cannabis exposure can result in lasting changes in brain structure, function, and behavior. [For a comprehensive review, see Navarrete et al. [13]].

CBD is often seen as safe because it does not have the same intoxicating effects as THC. However, like THC, CBD can cross the placenta and potentially harm a developing foetus [10]. Despite this, pregnant women use CBD as a natural remedy to relieve symptoms like nausea, insomnia, anxiety, and chronic pain [15]. While as much as 20% of pregnancies in the USA and Canada may involve the use of CBD-only products [4], the risks of using CBD during pregnancy are not fully understood. Recent studies in rodents suggest that prenatal CBD exposure (PCE) can have lasting effects on the offspring’s brain and behavior. These effects include anxiety, memory issues, and pain sensitivity changes in adulthood [16], [17]. Sex differences are also observed, with CBD affecting males and females differently. Thus, extended exposure of CD1 mice to CBD spanning from gestation through the first week after birth alters repetitive and hedonic behaviours in the adult progeny [18]. Swenson and colleges [17] also demonstrated that PCE influenced thermal pain sensitivity in a sex-specific manner. PCE produces sex-specific alterations in anxiety, memory and sensory gating during adolescence, associated with functional and molecular changes in prefrontal cortex and ventral hippocampus [19]. Furthermore, oral administration of CBD during gestation or lactation, or both, influenced pups’ survival, metabolism, obsessive compulsive- and anxiety-related behaviours and long-term object memory in a sex-dependent manner during adulthood [20]. Our previous research showed that exposing mice to a low dose of CBD in utero led to increased weight gain in male pups during the early stages of life. Additionally, the offspring of dams exposed to CBD during pregnancy exhibited sex-specific differences in communication, motor skills, and the ability to distinguish between things during early development [21]. Later in life, exposure to CBD during pregnancy caused sex-specific and region-specific changes in the brain’s electrical activity in the insular cortex, a deep brain region involved in processing sensory, emotional, and cognitive information [22]. Overall, these findings suggest that exposure to CBD in utero can have lasting effects on brain development and function.

Preclinical studies examining the neurodevelopmental effects of prenatal cannabidiol exposure (PCE) have largely focused on specific life stages, leaving gaps in our understanding of the full behavioral trajectory associated with PCE. To address this limitation, the present study takes a longitudinal approach to evaluate the neurodevelopmental trajectory following PCE in offspring of both sexes. The findings reveal sex-specific disruptions in developmental trajectories and vulnerabilities associated with gestational CBD exposure, challenging the notion that CBD is universally safe.

## Materials and methods

### Animals

Animals were treated in compliance with the European Communities Council Directive (86/609/EEC) and the United States NIH Guide for the Care and Use of Laboratory Animals. The French Ethical committee authorized the project APAFIS#18476-2019022510121076 v3. Adult male and female C57BL6/J (12–17 weeks age) were purchased from Charles River and housed in standard wire-topped Plexiglas cages (42 × 27 x 14 cm), in a temperature and humidity-controlled condition (i.e., temperature 21 ± 1 °C, 60 ± 10% relative humidity and 12 h light/dark cycles). Food and water were available ad *libitum*. Following a one-week acclimation period, female pairs were introduced to a single male mouse in the late afternoon. The day when a vaginal plug was observed was considered as day 0 of gestation (GD0), and pregnant mice were individually housed thereafter. Starting from GD5 until GD18, the pregnant dams received daily subcutaneous injections (s.c.) of either a vehicle or 3 mg/kg of CBD (obtained from the Nida Drug Supply Program). The CBD was dissolved in a vehicle solution composed of Cremophor EL (from Sigma-Aldrich), ethanol, and saline, with ratios of 1:1:18, respectively, and administered at a volume of 4 mL/kg. Control dams (referred to as “Sham”) received an equivalent volume of the vehicle solution. Upon each litter’s birth, the day was designated as postnatal day (PND) 0. Pups were weaned on postnatal day (PND) 21 and subsequently housed separately by sex. The experiments were carried out on the male and female offspring during early adolescence (PNDs 22-30) and adulthood (PNDs 90-120) (Figure 1). The exact sample size (n) for each experimental group/condition is indicated in the figure legends.

**Figure 1.**
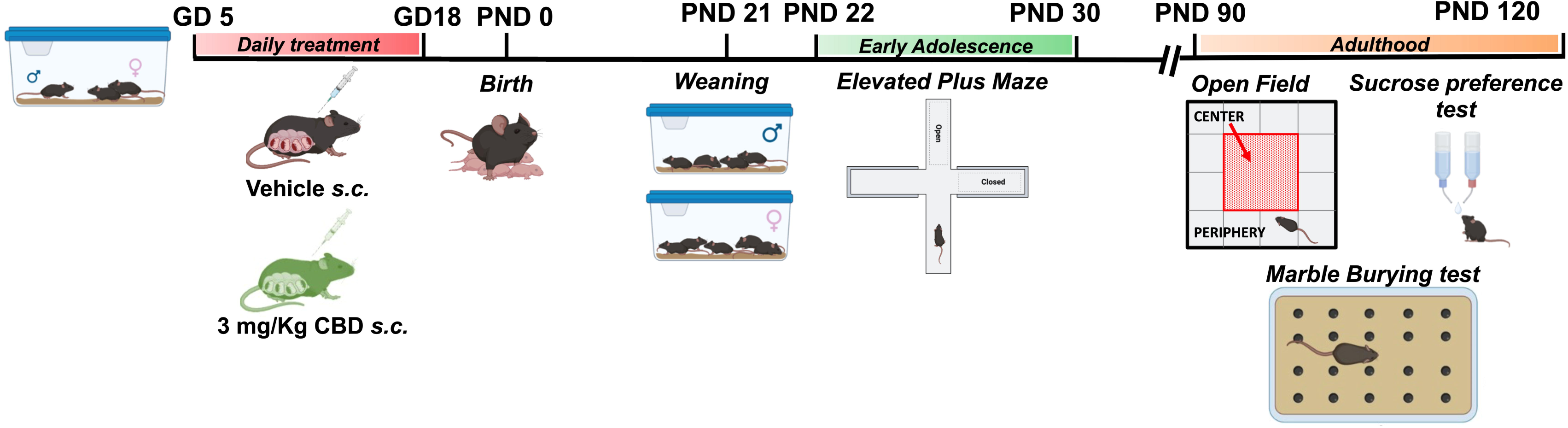
Experimental Timeline for Gestational Cannabidiol (CBD) Exposure and Behavioral Assessment. Mice were bred and treated with CBD once daily from gestational day (GD) 5 to GD 18. The date of birth was defined as post-natal day (PND) 0. Pups were weaned at PND 21 and housed separately by sex. During early adolescence, both sham and CBD-exposed offspring underwent the elevated plus maze test. Later in adulthood, both sexes were assessed using the open field, sucrose preference, and marble burying tests. Created with Biorender.com.

### Behavioral test

#### Elevated Plus-Maze

The elevated plus-maze apparatus comprised two open (50 × 10 × 40 cm) and two closed arms (50 × 10 × 40 cm) that extended from a common central platform (10 × 10 cm). Mice were individually placed on the central platform of the maze for 5 min. Each 5 min session was recorded with a camera positioned above the apparatus for subsequent behavioral analysis carried out using the Observer 3.0 software (Noldus, The Netherlands). The following parameters were analyzed:

- locomotory parameters: distance moved (cm), velocity (cm/s), the time spent moving and not (sec);
- time spent in the open arms: seconds spent on the open arms of the maze;
- time spent in the closed arms and in the center: seconds spent on the closed arms of the maze + seconds spent on the center of the maze;
- preference of open arms: (seconds spent on the open arms of the maze/total time spent in the maze-time spent on the open arms) x100.
- body state: time spent in normal, contracted and stretched state (sec).

#### Open field

The apparatus consisted of a Plexiglas arena 45 × 45 cm, illuminated by fluorescent bulbs at a height of 2 m above the floor of the open field apparatus (light intensity of 30 Lux). The floor was cleaned between each trial to avoid olfactory clues. Each animal was transferred to the open-field facing a corner and was allowed to freely explore the experimental area for 20 min. A video tracking system (Ethovision XT, Noldus Information Technology) recorded the exact track of each mouse as well as total distance traveled (cm), velocity (cm/sec), the total time spent moving and not (sec), the time spent (sec) and the entries in the corner defined as “periphery” and in the center area, the thigmotaxis (%) calculated as the (time spent in the periphery / time spent in the periphery + time spent in the center) x 100 [23], and the body state (normal, contracted and stretched state (sec)).

#### Sucrose Preference Test

Mice were tested for preference of a 2% sucrose solution, using a two-bottle choice procedure with slight modifications. Subjects were housed singly for the 3 days of test. Mice were given two bottles, one of sucrose (2%) and one of tap water. Every 24 h the amount of sucrose and water consumed was evaluated. To prevent potential location preference of drinking, the position of the bottles was changed every 24 h. The sucrose consumption was calculated as the amount of sucrose consumed in 24hours/mouse weight (g/Kg). While, the sucrose preference was calculated as the intake of sucrose solution consumed in 24hours/total fluid intake (%).

#### Marble Burying Test

20 black marbles were arranged in a fresh cage filled with 4–5 cm of bedding material. Mice were placed in one corner of the cage to start the trial, and the observation period was 20 min. Subsequently, the number of covered marbles (>2/3 of the marble surface covered by bedding material) was scored.

#### Statistical analysis

Statistics, graphs and figure layouts were generated with GraphPad Prism 10. Datasets were tested for the normality (D’Agostino-Pearson and Shapiro-Wilk) and outliers (ROUT test) before running parametric tests. Statistical significance of difference between means was assessed with two- or three-way ANOVA followed by Sidak’s multiple comparison post hoc tests as indicated in figure legends. When achieved, the significance was expressed as exactly *p*-value in the figures. The experimental results are described qualitatively in the main text, whereas experimental and statistical details, including sample size (N = number of animal), statistical test description and *p*-value are reported in Figure legends. Quantitative data are presented as Box and whisker plots, reporting median, min and max values, and superimposed scatter plots to show individual data points.

## Results

### Anxiety-related behavior during early adolescence in female progeny expose in-utero to CBD

Children exposed to cannabis in utero exhibit significantly higher rates of anxiety-related issues [24]. Similarly, gestational exposure to THC [25] or high dose of CBD (30 mg/Kg) [19] increases anxiety-related behavior in rats in adulthood and adolescence. In our previous study, we found that a low dose of CBD (3 mg/kg, GD5–GD18) during gestation caused sex-specific behavioral changes in neonatal pups [21]. Using the same PCE protocol in this study, we investigated whether early-life exposure leads to anxiogenic-like behaviors.

The elevated plus maze, a well-established method for evaluating anxiety responses in rodents [26], was used to assess anxiety behavior during early adolescence (PND 22–30, Figure 2). This method relies on the conflict between the closed and open arms of the maze. We quantified the time spent in each arm (Figure 2C–E) and found that gestational CBD exposure significantly increased the time CBD-exposed females spent in the closed arms and center of the maze (Figure 2C). Consequently, these females spent significantly less time in the open arms compared to Sham females, indicating a reduced preference for the open arms (Figure 2D, E). No differences in general locomotor activity were observed between Sham and CBD-exposed offspring, regardless of sex (Supplementary Figure 1). Specifically, distance traveled, speed, and time spent moving were similar across groups and sexes during the 5-minute test (Supplementary Figure 1B–E). Overall, these findings suggest that gestational CBD exposure induces a sex-specific anxiogenic effect in female mice, which was not observed in males.

**Figure 2.**
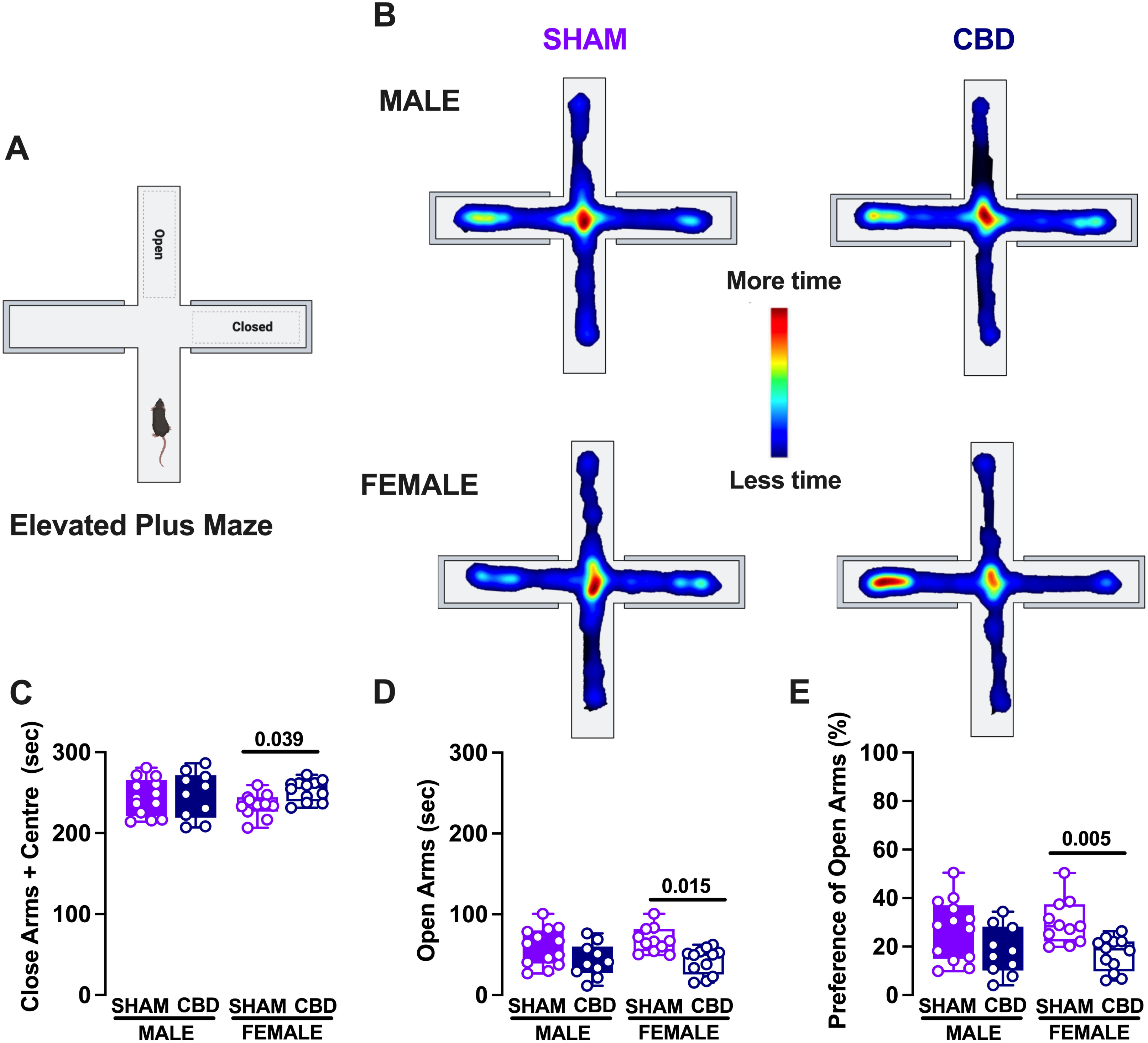
Anxiety-related behavior during early adolescence is selectively increased in female progeny exposed to gestational CBD. (A) Schematic representation of the Elevated Plus Maze apparatus highlighting the open and closed arms. (B) Heatmap presentation of the time spent in each arm of the maze over 300 seconds. Hot colors represent areas visited most frequently, and cold colors represent areas visited least frequently. (C) Female offspring exposed to CBD spent more time in the closed arms and center of the maze compared to the control group. In contrast, this behavior was similar in male offspring across both groups. (D, E) CBD-exposed females spent less time in the open arms, showing a lower preference for them compared to control females. (C-E) Data are represented as box and whisker plots (minimum, maximum, median). Individual data points represent one animal. Statistical analysis was performed using two-way ANOVA followed by Šídák’s multiple comparison test. *P-values < 0.05 are depicted in the graph. The sample sizes for each group were: SHAM MALE (N = 13 dark violet), CBD MALE (N = 10 dark blue), SHAM FEMALE (N = 11 light violet), CBD FEMALE (N = 12 light blue).

### The risk assessment behavior during the EPM task was enhanced in a sex-specific manner by in utero CBD exposure

Stretch-attend postures (SAP), where a rodent lowers its back, elongates its body, and either stands still or moves forward very slowly [27], are generally associated with anxiety. This posture represents a type of risk-assessment behavior thought to reflect an approach-avoidance tendency [28].

To determine whether the anxiety phenotype observed in the EPM task was linked to changes in risk-assessment behavior, we performed a detailed analysis of body posture during the 5-minute test (Figure 3). We quantified the total time spent in normal, stretched-attend, and contracted postures among Sham and CBD-exposed offspring of both sexes (Figure 3A–C). Our results showed that CBD-exposed females spent significantly less time in normal body posture compared to Sham females (Figure 3A). Importantly, these females spent more time in the stretched-attend posture than both Sham females and CBD-exposed males (Figure 3C). No significant differences were found in the time spent in a contracted posture across groups (Figure 3B).

**Figure 3.**
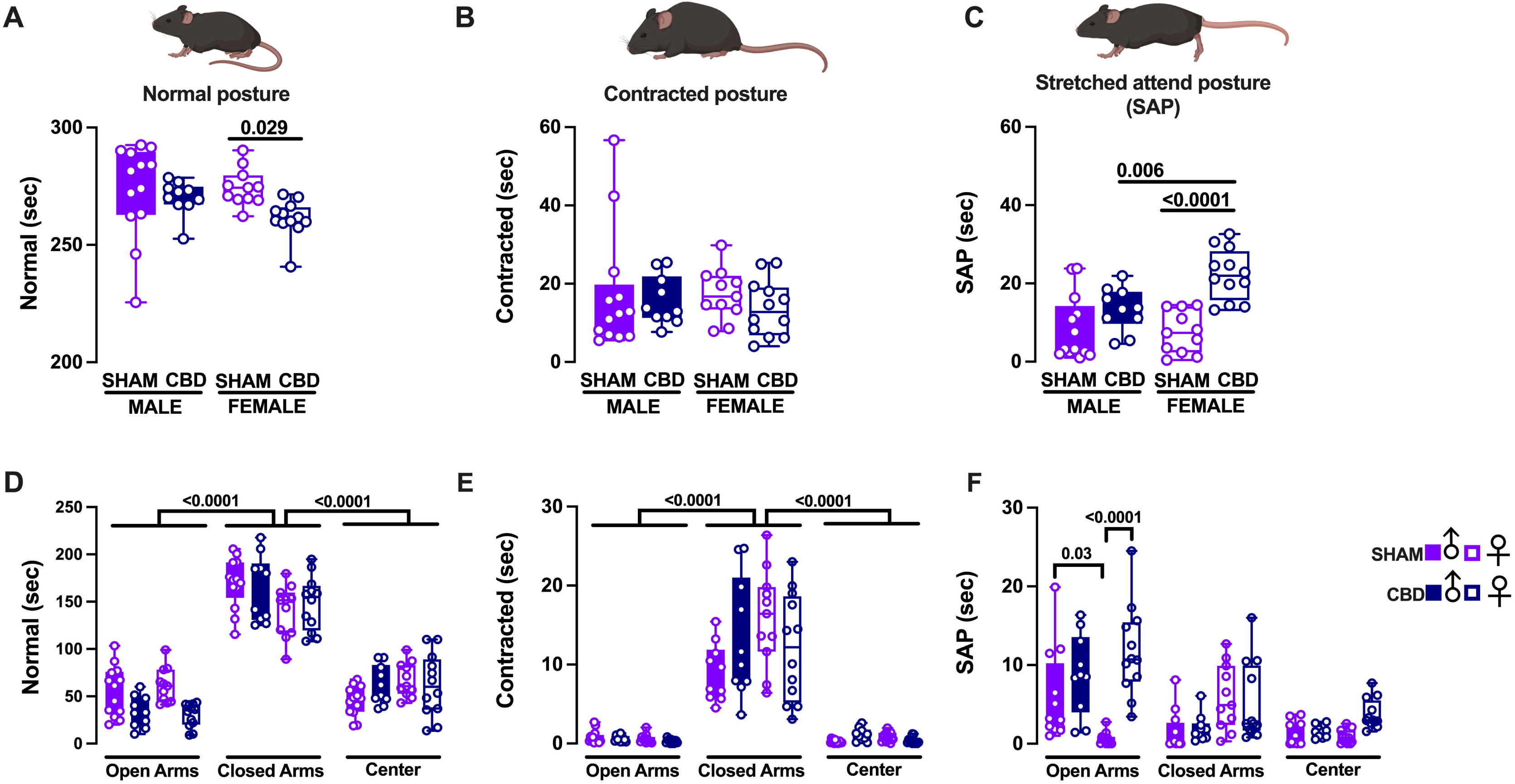
In utero CBD exposure increases risk assessment in female offspring. (A) Qualitative (top) and quantitative (bottom) analysis of body posture during the EPM task revealed that gestational CBD exposure reduced the time spent in a normal body posture in female offspring compared to Sham females. (B) A similar analysis of the contracted posture showed no significant differences in the time spent in this posture based on treatment or sex. (C) In contrast, the analysis of the stretched attend posture (SAP) revealed that female CBD-exposed animals spent more time in SAP compared to both sham females and CBD-exposed males. (D, E) Quantitative analysis of normal, contracted, and SAP in each arm of the EPM indicated that, regardless of sex or treatment, both sham and CBD-exposed offspring spent significantly more time in normal and contracted postures in the closed arms of the maze compared to the open and center arms. (F) Sham males spent more time in the SAP in the open arms of the maze compared to Sham females. After CBD gestational exposure, both male and female progeny spent similar amounts of time in SAP in the open arms, although CBD-exposed females remained in this posture longer than their Sham counterparts. Across different maze arms, Sham males spent a consistent amount of time in SAP, while Sham females spent less time in this posture in the open arms compared to the closed arms. Lastly, CBD-exposed females spent significantly more time in the stretched attend posture (SAP) in the open arms compared to the closed and center arms of the maze. (A-F) Data are represented as box and whisker plots (minimum, maximum, median). Individual data points represent one animal. Statistical analysis was performed using two-way or three-way ANOVA, followed by Šídák’s multiple comparison test for panels A-C and D-F, respectively. *P-values < 0.05 are indicated in the graph. The sample sizes for each group were as follows: SHAM MALE (N = 13, dark violet), CBD MALE (N = 10, dark blue), SHAM FEMALE (N = 11, light violet), and CBD FEMALE (N = 12, light blue).

To further explore correlations between body posture and maze behavior, we analyzed body states within each arm of the maze separately (Figure 3D–F). Both Sham and CBD-exposed animals, regardless of sex, spent more time in normal and contracted postures in the closed arms compared to the open and center arms (Figure 3D, E). Among Sham offspring, males spent more time in the stretched-attend posture in the open arms than females (Figure 3F). However, following in utero CBD exposure, male and female progeny spent a comparable amount of time in this posture in the open arms (Figure 3F). Notably, CBD-exposed females spent significantly more time in the stretched-attend posture in the open arms compared to their Sham counterparts (Figure 3F).

Additionally, Sham males maintained a consistent amount of time in the stretched-attend posture across all maze arms, while Sham females spent less time in this posture in the open arms compared to the closed arms (Figure 3F). In contrast, CBD-exposed females spent significantly more time in the stretched-attend posture in the open arms than in the closed and center arms (Figure 3F). These findings reinforce the anxious phenotype observed during the EPM test and emphasize the sex-specific anxiogenic effects of prenatal cannabinoid exposure.

### The anxiety-related phenotype persists into adulthood in CBD-exposed females

The developmental trajectory of anxiety typically follows an inverse relationship with the maturation of emotion regulation, with anxiety increasing during adolescence and decreasing as individuals transition into young adulthood [29] [30]. To examine the time course of anxiety, we conducted the open field task during adulthood (Figure 4, Supplementary Figure 2). A sex-specific effect of gestational CBD exposure was observed in the time spent in different compartments of the arena. Specifically, CBD-exposed females, but not males, spent significantly more time in the periphery and less time in the center compared to both Sham females and CBD-exposed males (Figure 4C, D). Additionally, prenatal cannabinoid exposure (PCE) induced a sex-specific anxiogenic effect, increasing thigmotaxis exclusively in CBD-exposed females relative to Sham females and CBD- exposed males (Figure 4E).

**Figure 4.**
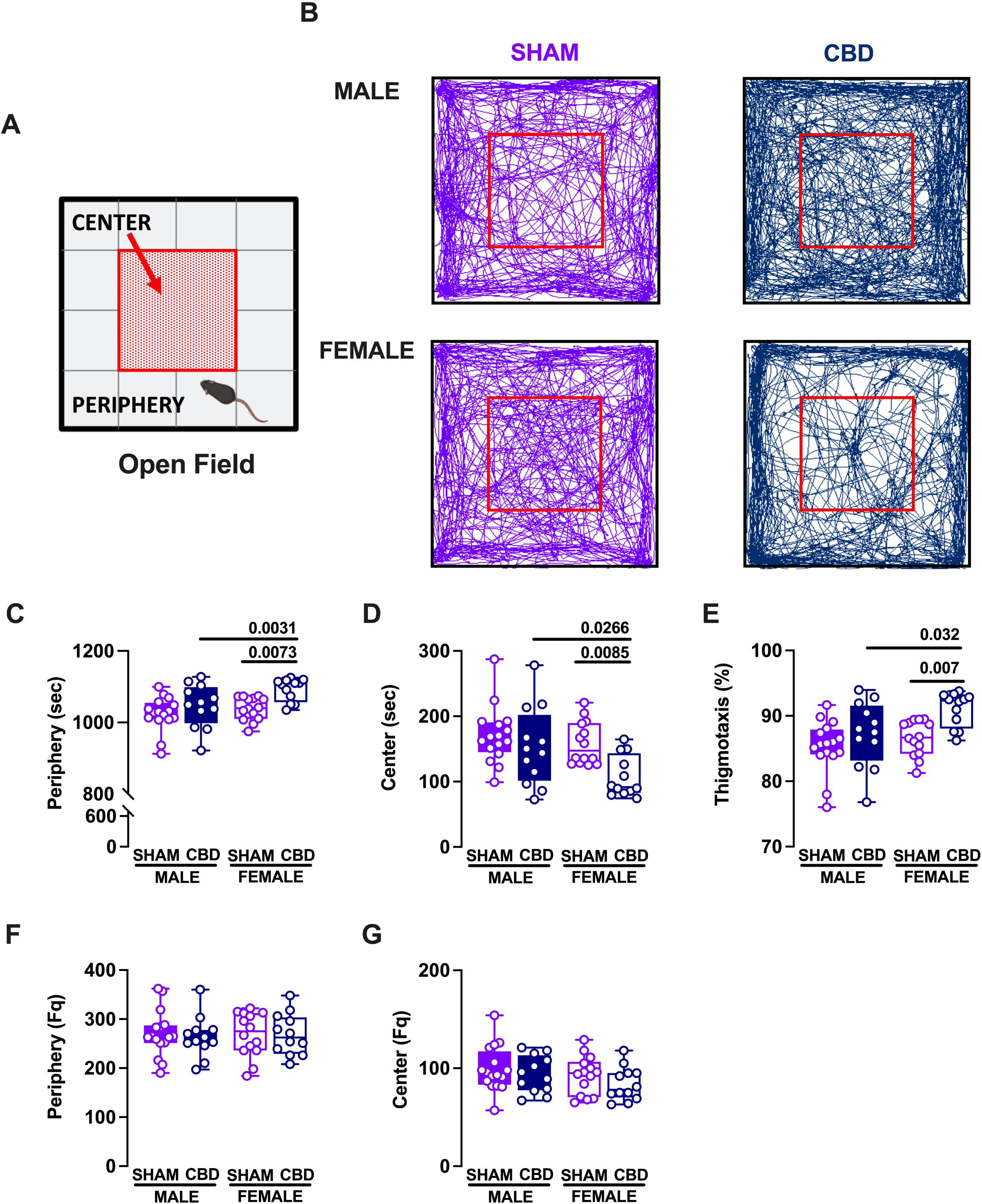
Persistent anxiety phenotype in adulthood results from prenatal exposure to CBD. (A) The schematic illustration shows the Open Field apparatus, with the central square labeled as “center” (red arrow) and the surrounding area labeled as “periphery.” (B) Representative track traces of mouse movement during a 20-minute test session are shown for Sham (violet, left) and CBD-exposed (blue, right) adult mice of both sexes. Following in-utero CBD exposure, adult CBD females spent significantly more time in the periphery (C) and less time in the center (D) compared to Sham females and CBD-exposed males. (D) The level of thigmotaxis, expressed as the percentage of time spent in the periphery, was greater in CBD-exposed females compared to both Sham females and CBD-exposed males. (F, G) In contrast, gestational CBD exposure did not affect the number of entries into the periphery or center areas of the maze. (C-G) Data are represented as box and whisker plots (minimum, maximum, median), with individual data points representing one animal. Statistical analysis was performed using two-way ANOVA followed by Šídák’s multiple comparison test, with *P-values < 0.05 indicated on the graph. The sample sizes for each group were as follows: SHAM MALE: N = 16; CBD MALE: N = 12; SHAM FEMALE: N = 14; CBD FEMALE: N = 12.

Despite these differences, entry rates into the periphery and center of the arena were similar across Sham and CBD-exposed groups, regardless of sex (Figure 4F, G). We also analyzed time spent in different body postures (Figure 5). Consistent with earlier findings from adolescence, CBD-exposed females spent less time in a normal body posture compared to Sham females and CBD-exposed males (Figure 5A). The total time spent in a contracted posture was similar across all treatment groups and sexes (Figure 5B). Interestingly, among Sham offspring, females spent significantly more time in the stretch-attend posture (SAP) compared to males. However, after prenatal cannabinoid exposure, this sex difference disappeared, as CBD-exposed females spent significantly more time in SAP than both Sham females and CBD-exposed males (Figure 5C).

**Figure 5.**
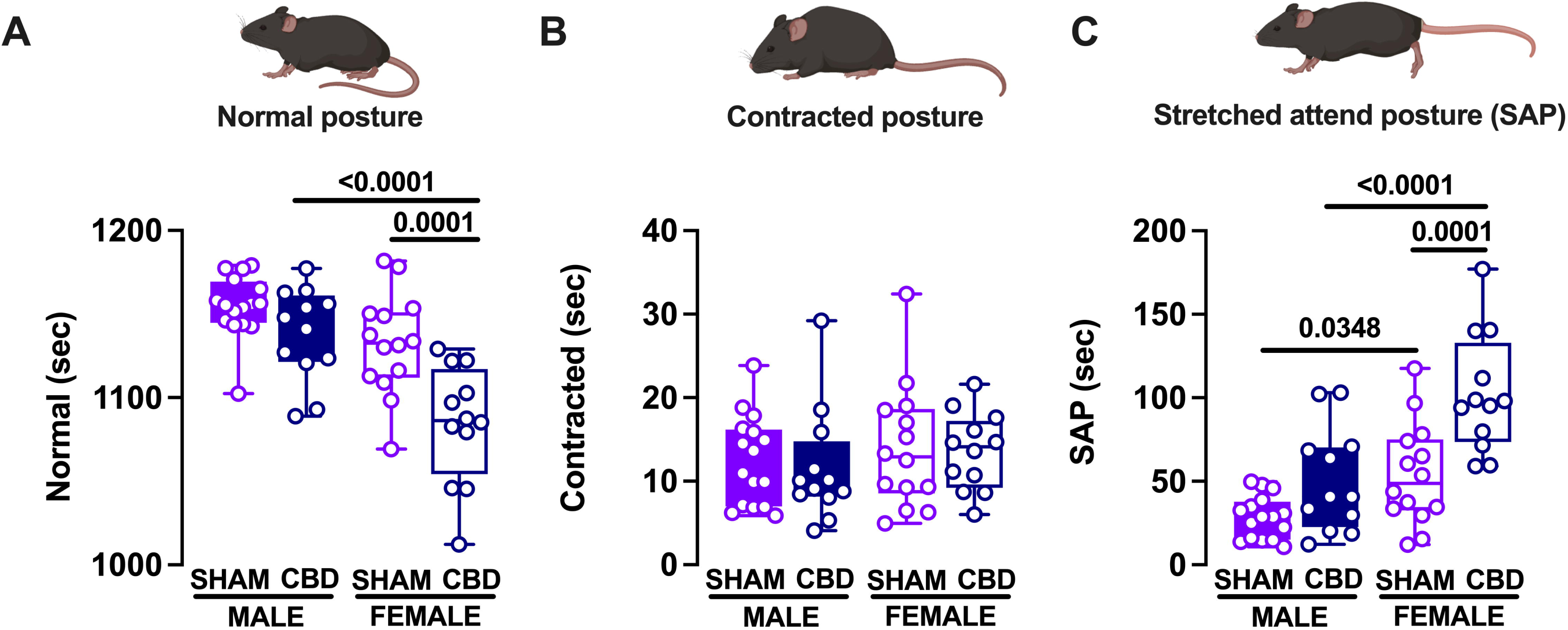
Risk assessment behavior is selectively enhanced in females during adulthood following in utero CBD exposure. (A) Qualitative (top) and corresponding quantitative (bottom) representations of normal body posture during the open field task show that CBD-exposed females spent significantly less time in the normal body state compared to their Sham counterparts and CBD-exposed males. (B) A similar analysis of contracted posture revealed that the total time spent in a contracted body state was similar between Sham and CBD-exposed animals in both sexes. (C) Analysis of the SAP indicated that both Sham and CBD-exposed female progeny spent more time in the SAP. However, CBD-exposed females spent significantly more time in this state compared to Sham females, while no significant differences were observed in males. (A-C) Data are represented as box and whisker plots (minimum, maximum, median), with individual data points representing one animal. Statistical analysis was performed using two-way ANOVA followed by Šídák’s multiple comparison test. *P-values < 0.05 are depicted in the graph. The sample sizes for each group were: SHAM MALE N = 16 (dark violet), CBD MALE N = 12 (dark blue), SHAM FEMALE N = 14 (light violet), CBD FEMALE N = 12 (light blue).

Finally, no differences in distance traveled, velocity, or mobility were observed during the 20-minute task between Sham and CBD-exposed offspring, regardless of sex (Supplementary Figure 2B–E). Overall, these findings indicate that the anxiety phenotype observed in CBD-exposed females during early adolescence persists into adulthood, following a sex-specific developmental trajectory.

### The responsivity of adult CBD-exposed progeny to natural reinforcing stimuli in the sucrose preference test was increased by fetal CBD exposure

Epidemiological studies have reported a significant positive association between anxiety levels and sugar intake [31]–[34].

To investigate whether the persistent anxiety phenotype observed in CBD-exposed females during early and late development influenced their sugar consumption, we conducted a sucrose preference test on both Sham and CBD-exposed offspring (Figure 6 and Supplementary Figure 3).

**Figure 6.**
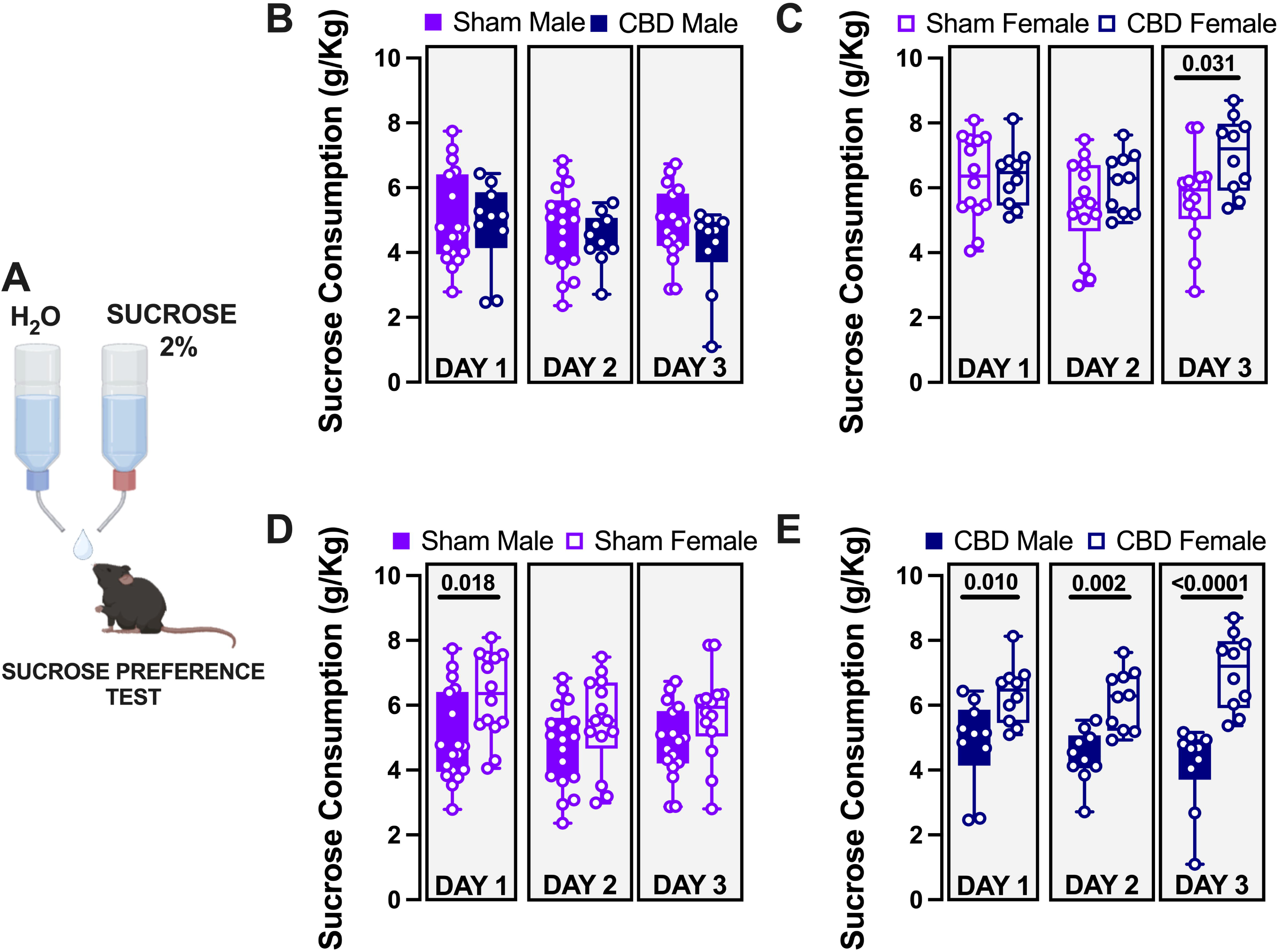
Fetal CBD increased the responsivity of adult female progeny to natural reinforcing stimuli in the sucrose preference test. (A) Schematic representation of the sucrose preference test design: mice were individually housed in single cages with continuous access to both water and a 2% sucrose solution for 3 days. (B) Throughout the duration of the test, male animals exposed to CBD consumed comparable amounts of sucrose as Sham animals. (C) In contrast, on the third day, CBD-exposed females showed higher sucrose consumption compared to Sham females. (D) Among Sham animals, females consumed more sucrose than males on the first day of the test, but consumption was similar between sexes on subsequent days. (E) Gestational CBD exposure increased sucrose consumption only in CBD-exposed females compared to CBD-exposed males. (B-H) Data are represented as box and whisker plots (minimum, maximum, median). Individual data points represent one animal. Statistical analysis was performed using two-way ANOVA followed by Šídák’s multiple comparison test. *P-values < 0.05 are depicted in the graph. The sample sizes for each group were: SHAM MALE (N = 18, dark violet), CBD MALE (N = 10, dark blue), SHAM FEMALE (N = 14, light violet), CBD FEMALE (N = 10, light blue).

Our analysis revealed sex-specific differences in sucrose consumption between the treatment groups. Male animals exposed to CBD consumed similar amounts of sucrose compared to their Sham counterparts (Figure 6B). However, on the third day, CBD-exposed females demonstrated higher sucrose consumption than Sham females (Figure 6C). Sucrose consumption was comparable between males and females in the Sham group (Figure 6D). In contrast, within the CBD-exposed group, females consistently consumed significantly more sucrose than males throughout the testing period (Figure 6E).

To ensure that isolation during the testing period did not negatively affect body weight (BW), we monitored BW over the three days. As expected, males weighed significantly more than females in both Sham and CBD-exposed groups, and no changes in BW were observed during the testing period (Supplementary Figure 3A). Additionally, no differences in sucrose preference were detected across treatment groups or sexes (Supplementary Figure 3B-E).

These findings are consistent with clinical data, suggesting a potential link between anxiety traits and increased sucrose consumption.

### Repetitive behaviors in the marble burying task of adult offspring were increased by gestational CBD exposure

The marble burying test was employed to evaluate anxiety-like and/or repetitive behaviors in our mouse groups [35]–[37] (Figure 7). The test revealed that CBD-exposed offspring buried significantly more marbles compared to the Sham group, irrespective of sex (Figure 7B). In contrast, Sham progeny of both sexes buried a similar number of marbles during the test (Figure 7B). These findings suggest that in utero CBD exposure in mice significantly influences both anxiety-related and repetitive behaviors.

**Figure 7.**
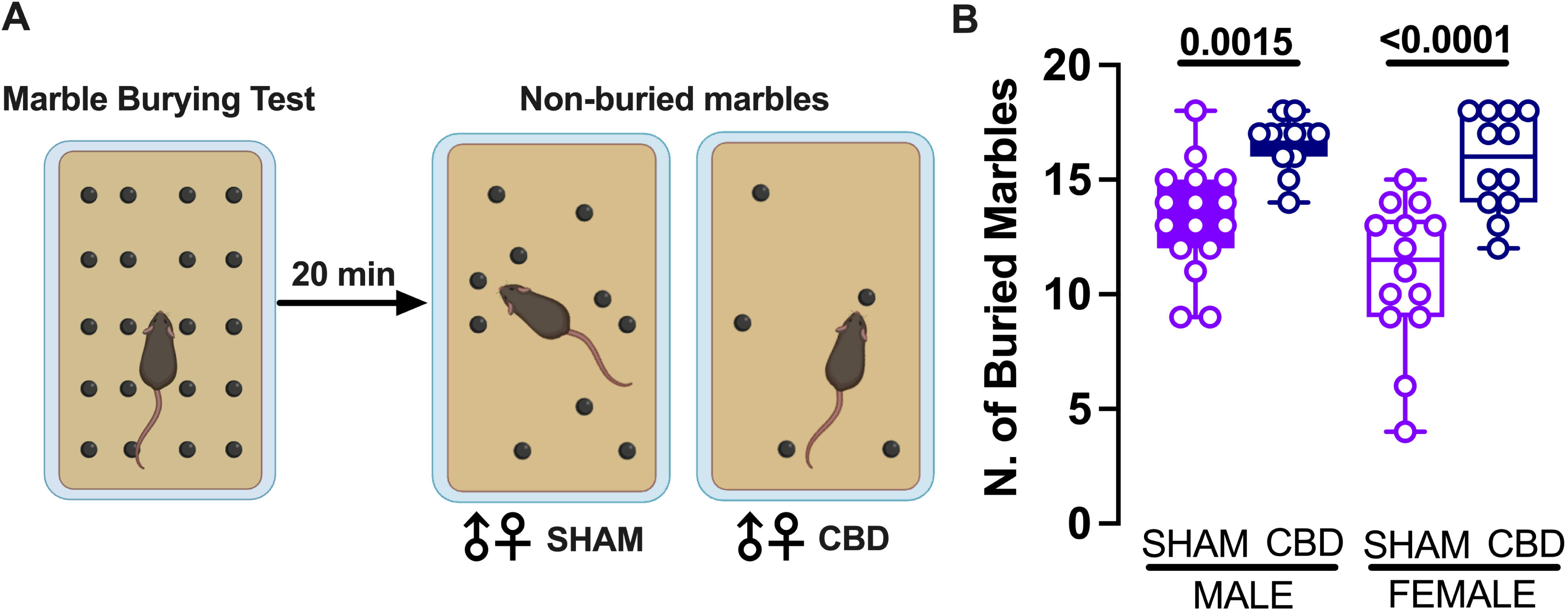
Repetitive behaviors in the marble burying task were increased in adult offspring exposed to gestational CBD. (A) The marble burying paradigm (left) showing representative results for Sham and CBD animals of both sexes (right). (B) Both male and female offspring prenatally exposed to CBD buried significantly more marbles compared to their Sham counterparts. (C) Data are represented as box and whisker plots (minimum, maximum, median). Individual data points represent one animal. Statistical analysis was performed using two-way ANOVA followed by Šídák’s multiple comparison test. *P-values < 0.05 are indicated in the graph. The sample sizes for each group were: SHAM MALE (N = 16, dark violet), CBD MALE (N = 12, dark blue), SHAM FEMALE (N = 14, light violet), CBD FEMALE (N = 12, light blue).

## Discussion

Exposure to environmental factors such as cannabis during critical developmental periods has been associated with an increased risk of neuropsychiatric disorders later in life [13], [38], [39]. In this longitudinal study, we investigated the effects of prenatal cannabis exposure (PCE) on postnatal neurodevelopment. Our findings reveal that gestational CBD exposure leads to sex-specific behavioral changes, with females—but not males—exhibiting an anxiety-like phenotype during early adolescence. This anxiety persists into adulthood and is accompanied by altered reward responsiveness. Additionally, adult offspring of both sexes exposed to CBD demonstrate increased repetitive behaviors.

These findings suggest that PCE creates sex-specific developmental vulnerabilities to neuropsychiatric conditions, particularly anxiety. This aligns with clinical evidence highlighting significant sex differences in neuropsychiatric disorders [40]–[42]. Anxiety, one of the most common mental health disorders, disproportionately affects women throughout their lives [43].

Maternal cannabis use has been associated with anxiety-related traits in early childhood and reduced placental gene expression [24], [38], [44]. Previous animal studies have also linked PCE to behavioral changes, including increased anxiogenic behaviors in offspring [45]. Extending these findings to CBD, a cannabis compound often perceived as safe, we found that maternal exposure to a low dose of CBD induces sex-specific anxiety behaviors. Female offspring spent less time in, and showed a lower preference for, the open arms of the elevated plus maze (EPM) during early adolescence.

This vulnerability may stem from differences in how CBD accumulates in the embryonic brain; females have been shown to retain higher CBD levels than males following exposure to the same dose [18]. Additionally, alterations in 5HT1A serotonin receptors, which are sex-specific, may contribute to this vulnerability due to the receptor’s role in regulating stress, anxiety, and depression [19], [46], [47]. Supporting this, our previous studies demonstrated greater impairments in sensory and cognitive development in female neonates exposed to CBD compared to males [21].

Our analysis of specific anxiety-related behaviors supports these findings. Female offspring exposed to CBD exhibited increased stretch-attend posture (SAP), a recognized risk-assessment behavior associated with anxiety in rodent models [27], [48]–[55]. These behaviors persisted into adulthood, where CBD-exposed females showed heightened thigmotaxis (the tendency to stay near walls) during the open field task, a key indicator of anxiety [56].

In addition to anxiety traits, gestational CBD exposure caused sex-specific changes in reward responsiveness. Adult CBD-exposed females consumed more sucrose than their male counterparts, suggesting altered reward processing. Although limited, some epidemiological studies report a positive correlation between anxiety and sugar intake, hinting at a potential comorbidity [57]–[59]. We also found that PCE increased repetitive behaviors in both sexes during adulthood, as evidenced by performance in the marble burying test. This task is commonly used to assess anxiety, obsessive-compulsive tendencies, or repetitive actions in rodents [36], [37]. Obsessive-compulsive disorder (OCD), a developmental neuropsychiatric illness with early (adolescence) or late (young adulthood) onset, often co-occurs with anxiety disorders [60]–[63]. Our findings suggest that PCE increases heterogeneous risks for psychiatric conditions across the lifespan.

Mechanistic insights reveal that gestational CBD exposure disrupts neuronal development in the insular cortex (IC) into adulthood. The IC is a critical integration hub connecting sensory, emotional, motivational, and cognitive systems [64]–[66]. Dysfunction in the IC has been strongly linked to the onset of psychiatric conditions, including anxiety and OCD [67]–[71]. These disruptions likely underlie the observed sex-specific trajectories of anxiety and repetitive behaviors following PCE.

In conclusion, this longitudinal study provides robust evidence that gestational CBD exposure induces sex-specific disruptions in neurodevelopmental trajectories. These effects emerge early in utero and persist throughout life, increasing the risk for neuropsychiatric disorders such as anxiety and OCD. Our findings underscore the need for caution in using CBD during pregnancy, given its potential long-term impact on offspring mental health.

## Author Contributions

D.I.: conceptualization, data curation, formal analysis, validation, writing— original draft, review and editing. A.C.-R.: formal analysis. P.C.: conceptualization, supervision. O.J.J.M.: conceptualization, supervision, funding acquisition, methodology, project administration, writing—original draft, review, and editing. All authors have read and agreed to the published version of the manuscript.

## Acknowledgements

The authors are grateful to the Chavis-Manzoni team members for helpful discussions.

## Figure legends

**Supplementary Figure 1. The general locomotory profile of CBD progeny during early adolescence is not modified by gestational CBD exposure.** (A) Schematic representation of the EPM apparatus, highlighting the open and closed arms. (B-E) Gestational CBD exposure did not modify the distance moved, velocity, or total mobility time of CBD progeny of both sexes. (B-E) Data are represented as box and whisker plots (minimum, maximum, median). Individual data points represent one animal. Statistical analysis was performed using two-way ANOVA followed by Šídák’s multiple comparison test. *P-values < 0.05 are indicated in the graph. The sample sizes for each group were: SHAM MALE (dark violet, N = 13), CBD MALE (dark blue, N = 10), SHAM FEMALE (light violet, N = 11), CBD FEMALE (light blue, N = 12).

**Supplementary Figure 2. The locomotory parameters during adulthood were not altered by in-utero CBD exposure.** (A) Schematic representation of the Open Field apparatus. The central square is labelled as “center” (red arrow), while the surrounding area is labelled as “periphery.” (B-E) Gestational CBD exposure did not modify the distance moved, velocity, or total mobility time of CBD progeny in both sexes. (B-E) Data are represented as box and whisker plots (minimum, maximum, median). Individual data points represent one animal. Statistical analysis was performed using two-way ANOVA followed by Šídák’s multiple comparison test. *P-values < 0.05 are indicated in the graph. The sample sizes for each group were: SHAM MALE N = 16 (dark violet), CBD MALE N = 12 (dark blue), SHAM FEMALE N = 14 (light violet), CBD FEMALE N = 12 (light blue).

**Supplementary Figure 3. Sucrose preference was compared across treatment and sex.** (A) No differences in body weight were observed between Sham and CBD-exposed animals of both sexes during the 3-day test. (B-E) Gestational CBD exposure did not affect sucrose preference in CBD-exposed animals compared to Sham animals across sex and treatment. (B-E) Data are represented as box and whisker plots (minimum, maximum, median). Individual data points represent one animal. Statistical analysis was performed using two-way ANOVA followed by Šídák’s multiple comparison test. *P-values < 0.05 are depicted in the graph. The sample sizes for each group were: SHAM MALE N = 18 (dark violet), CBD MALE N = 10 (dark blue), SHAM FEMALE N = 14 (light violet), CBD FEMALE N = 10 (light blue).

