## Supplementary figures and images for "Sex-specific disruptions in the developmental trajectory of anxiety due to prenatal cannabidiol exposure"

### Supplementary Figure 1

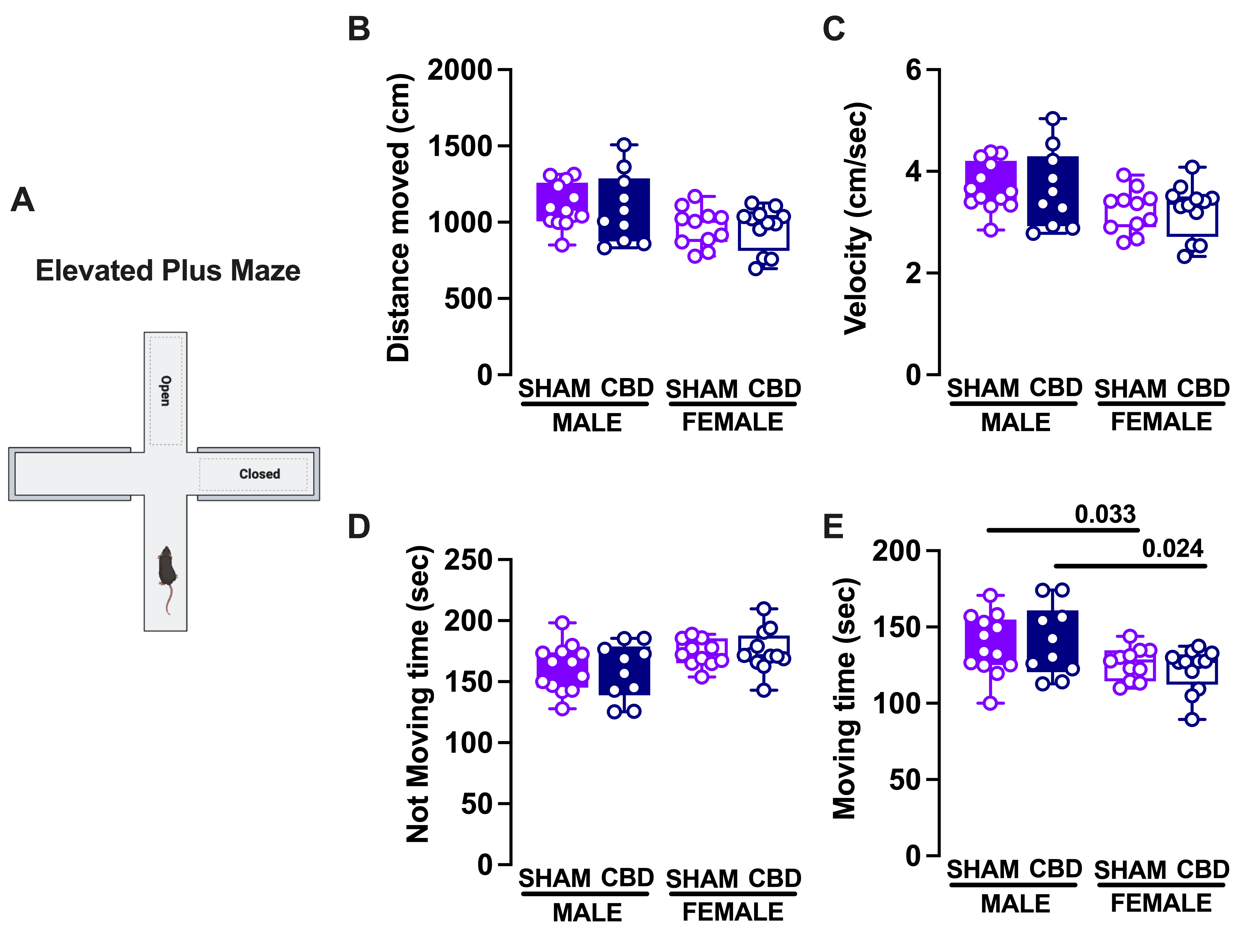

### Supplementary Figure 2

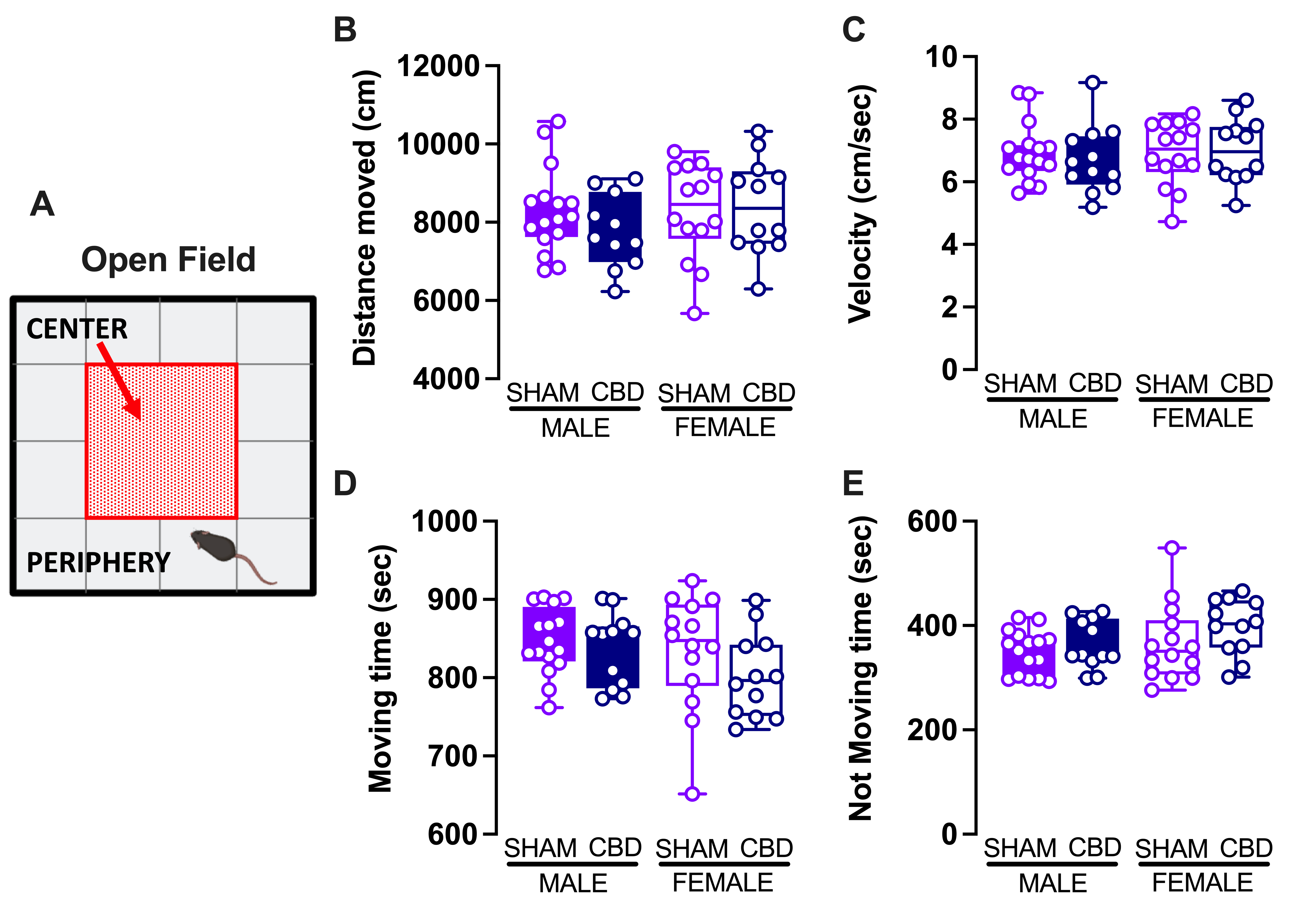

### Supplementary Figure 3

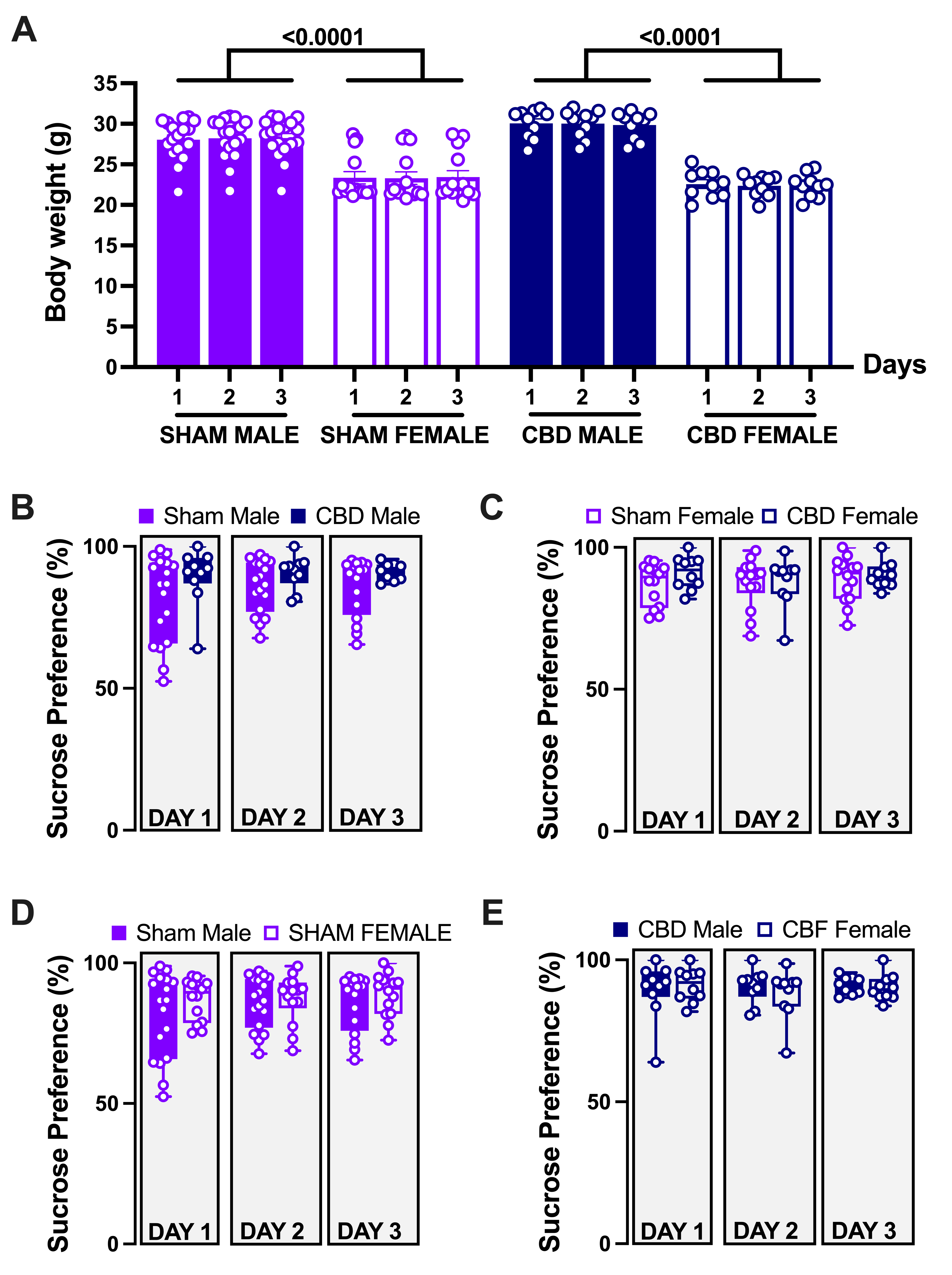
